# Evidence for a Male Bias in Atlantic Blue Crab Pot-Based Sampling

**DOI:** 10.1101/2023.05.09.538440

**Authors:** Jeff F. Brunson, Kimberly A. Sitta, Michael R. Kendrick, Peter R. Kingsley-Smith

## Abstract

The Atlantic Blue Crab *Callinectes sapidus* is an ecologically- and economically-important species that supports one of the highest valued commercial fisheries in coastal South Carolina, USA. Researchers at the South Carolina Department of Natural Resources conduct multiple surveys to monitor the status of the Atlantic Blue Crab using a variety of gear configurations. Demographic characterizations, however, can often be influenced by sampling gear-related biases. We compared blue crab sex ratios between passive, pot-based, sampling and active, trawl-based, sampling across three estuaries in the fall and for year-round sampling in a single estuary. For fall sampling, the percent of males in pots was 20.1% higher than values observed for trawl-based sampling, while an overall male bias of 22.7% was observed for year-round sampling; however, this bias was only significant in certain months. Our findings suggest that while different sampling gears may offer their own advantages, such as greater suitability to effectively sample specific habitats, the resulting characterizations of population demographics can differ among gear types. Recognizing gear related biases is important for ensuring that field surveys are representative of populations, particularly when sex-specific modeling approaches are used as part of stock assessments to determine population status.

## Introduction

Fishery-independent surveys are widely used to assess the status of populations of ecologically- and economically-important finfish and invertebrate species (Gunderson 1993). These surveys provide information on population demographics and abundance trends that can be used to improve our understanding of life history and to inform species stock assessments (*e*.*g*., Rago 2005). Depending on the type of habitat and particular species of interest, fishery-independent survey methods can involve the use of either passive (*e*.*g*., traps, pots, gillnets) or active (*e*.*g*., electrofishing, vessel-based trawling) sampling gear (see Rago 2005 for review). Each gear type, however, has its own potential biases, such that careful considerations must be made when selecting which gear type to use when assessing a population in order to avoid inaccurate interpretations of the derived data related to abundance trends and life history parameters.

It is well documented that a particular gear type may select for certain species (Valentinsson and Ulmestrand 2008; Mehdi et al. 2021), sizes (Rudstam et al. 1984; Reid et al. 2007), ages (Ralph and Lipcius 2014), and sexes (Hubert et al. 2012). For example, Mehdi et al. (2021) examined the gear selectivity of minnow traps, Windermere traps, and electrofishing sampling gear and found that finfish community assemblages differed significantly among gear types. In addition, in the case of the European Spiny Lobster *Palinurus elephas*, trap-based sampling results in sex ratios that are biased towards females (Goñi et al. 2003). While active sampling is thought to be less selective than passive sampling (Reid et al. 2007; Mehdi et al. 2021), active sampling is often more costly and labor intensive than passive sampling (Rago 2005; Xu et al. 2015) and may be restricted to suitable habitat types (Rago 2005). Passive sampling gear can be deployed across a wide range of habitats and is often a lower cost option (Bellchambers and de Lestang 2005); however, its effectiveness largely depends on the morphology and behavior of the target species (Reid et al. 2007). For example, entanglement gear, such as gillnets, requires that the animal encounter the gear and have a specific body shape and size to be retained (Hubert et al. 2012).

A variety of gear types are used to sample crustacean species, with most surveys conducted using either pots (Abbe 2002; Bellchambers and de Lestang 2005) or bottom trawls (Eggleston et al. 2004; Chen et al. 2006; Miller et al. 2011; Sturdivant and Clark 2011). Baited pots are often chosen due to their versatility, but their effectiveness is strongly dependent upon the feeding behavior of the targeted species. The catch of crustaceans using pots is influenced by factors such as soak time, choice of bait, and pot design (Boutillier and Sloan 1987), as well as behavioral traits of the target species (Jury et al. 2001; Sturdivant and Clark 2011). For example, Taggart et al. (2004) found that crab pots were biased against ovigerous females of the Dungeness Crab *Cancer magister*, potentially due to their lower mobility and feeding rates. Crab pots may also select for certain sizes of crabs, with studies finding that immature and subadult crabs are less likely to be captured than larger adults (Williams and Hill 1982; Smith et al. 2004).

The Atlantic Blue Crab *Callinectes sapidus* Rathbun is an ecologically- and economically-important invertebrate species in South Carolina (hereafter SC), with annual commercial landings typically exceeding three million pounds and valuing approximately $5 million (ACCSP, 2022), while also supporting a substantial recreational fishery. Atlantic Blue Crabs also fulfil a number of ecosystem services, including maintaining salt marsh integrity (Altieri et al. 2012), creating trophic linkages between benthic and pelagic food webs (*e*.*g*., Able et al. 2018), and serving as important prey items at all life stages (Williams 1984; Hines et al. 1990). Effective monitoring of Atlantic Blue Crab may improve our understanding of their potential influences on the broader estuarine community and ecosystem function (Silliman and Bertness 2002).

Previous analyses of trawl data in SC have shown that the sex ratio of blue crabs in trawls can be female dominated (*e*.*g*., Archambault et al. 1990); however, Eldridge and Waltz (1977) determined that the pot-based commercial catch in the southern part of the state was predominantly comprised of male Atlantic Blue Crab. It is unclear whether this discrepancy in the sex composition was due to the habitat where the Atlantic Blue Crabs were collected, or selectivity based on the gear itself. While a previous study by Bellchambers and de Lestang (2005) found that crab pots were more selective for male than female Flower Crab *Portunus pelagicus*, compared to either trawls or seines, no published studies have been conducted to test for sex biases related to sampling gear for Atlantic Blue Crab.

The present study investigated the potential for gear-specific sex bias in catches of Atlantic Blue Crabs. The specific objectives of the present study were to: (1) test for differences in the sex ratio of Atlantic Blue Crabs caught in pot-based and trawl-based sampling surveys across multiple estuaries; and (2) test for differences in the sex ratio of Atlantic Blue Crabs caught in pot-based and trawl-based sampling surveys at different time points throughout the year within a single estuary. Data generated from this study will be evaluated in the context of their implications for population assessments.

## [A] Methods

### [B] Data sources

#### [C] Passive gear

Passive gear sampling for Atlantic Blue Crabs was conducted using commercially available crab pots, constructed from 38 mm PVC-coated wire mesh and measuring 61 cm x 61 cm x 46 cm. Pots had bait wells and parlors and were equipped with a single 22.5 cm x 11 cm entrance funnel on each of the four vertical sides. Pots used in this survey were not equipped with the 6.0 cm escape rings, intended to facilitate the escapement of undersized crabs, that are required in commercial pots in SC. As such, crabs smaller than would typically be captured in commercial pots were retained in our sampling. Pots were tethered to buoys with sinking line and weighted with ≥ 1.6 mm steel rebar to minimize gear movements in swiftly moving currents generated from tidal ranges in excess 1.5 m. Within each of three estuaries, sampling occurred in three of five randomly selected blocks on a given day. Within each block, five pots were set approximately 100 m apart along a single line parallel with the shore, generally near where water depth changes rapidly. Pots were baited with Atlantic Menhaden, *Brevoortia tyrannus*, and allowed to soak for approximately four hours.

#### [C] Active gear

Trawling at a single station in each system was conducted monthly in the Ashley River, but only conducted in March, April, August, and December in the Dawho and Ashepoo Rivers. Trawling was conducted using 6.1 m flat otter trawls with 2.54 cm stretch mesh (1.27 cm bar mesh) and tickler chains. Nets were towed for 15 minutes at each station at approximately 1.3 m sec^-1^ (2.5 knots) and were deployed and retrieved using a hydraulic winch. For both active and passive gear sampling, carapace width and sex were recorded for all live Atlantic Blue Crab collected, prior to crabs being returned to the water.

### [B] Analytical approaches

#### [C] Across estuary fall comparison

To determine whether there was a sex bias in the catch of Atlantic Blue Crabs across estuaries, catch data from pot-based sampling were compared to trawl-based sampling data collected in three estuaries (Figure 1). Using a paired approach, data collected from pot-based sampling in November and December between 1988 and 2021 were compared to trawl data collected in December during the same time period.

**Figure 1.**
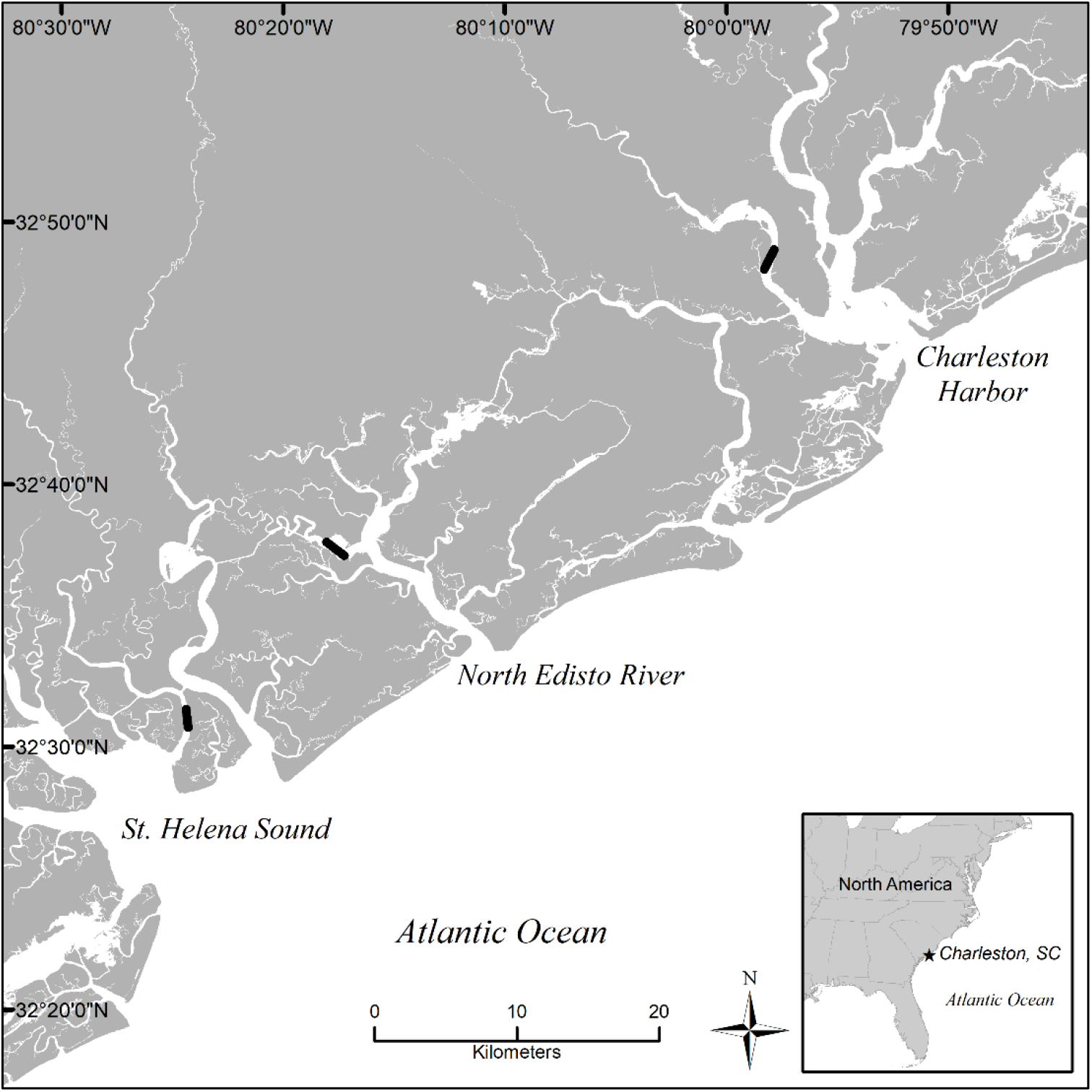
Sampling locations used for comparative analyses of catches of blue crabs from trawl surveys and commercial-style crab pots. Bold lines indicate segments of waterbodies sampled in pot-based and trawl-based surveys.

#### [C] Within estuary year-round comparisons

To determine if there was a sex bias in the catch of crabs within an estuary throughout the year, data from pot-based sampling were compared to data from trawl-based sampling using a paired approach for year-round sampling in the Ashley River between 2003 and 2019.

### [B] Data

All Atlantic Blue Crab ≥ 80 mm CW were included in our analyses, as this was the minimum size that was consistently retained in the pots. For each pot sampling event, data were pooled across all pots to determine the male sex ratio as the percent of the total crab catch. Likewise, males as a percent of the total catch was calculated for each trawl. In cases where no crabs were collected in either survey during a particular month, that sampling pair was excluded from analyses. When more than one pot sampling event occurred at a specific location in November or December of a given year, or within the same month in the Ashley year-round sampling, only data from the sampling event closest in time to the trawl sampling event of that year were included in analyses.

#### [C] Interval between sampling events

To determine an appropriate period of time between paired pot- and trawl-based sampling events for later analysis, analyses using methods described in the *Statistical analysis* section below were conducted to determine if the number of days between paired sampling events (from five to 14 days between sampling events in fall sampling and from one to 14 days between for year round sampling events) resulted in statistically significant differences in sex ratios of catches between the two gear types. Based on the results of those analyses and to provide sufficient sample size for analysis of estuary specific data in the fall, as well as temporal trends in year-round sampling, 14 days was selected as the maximum number of days between paired sampling events for inclusion in subsequent analyses. Filtering the data in this way limited data available for analyses to those collected from 1990-2021 for the fall sampling across estuaries but did not affect the time-series of data collected year-round in the Ashley River.

### [B] Statistical analysis

Male crabs as a percent of the total catch in pot-based sampling was compared to the percent of male crabs in the corresponding trawl sampling event to determine if there was a sex bias in pot-based sampling relative to trawl sampling using a paired, two-tailed Wilcoxon Signed Rank test. All statistical analyses were conducted in R v. 4.1.2 (R Core Team 2021) using the wilcox.exact function to account for zeros and ties in the paired data using the “exactRankTests” package (Hothorn and Hornik 2022). Statistical significance was set at an α value of 0.05.

## [A] Results

### [B] Across estuary fall comparisons

For data from fall sampling, the percent of male crabs in pots was significantly higher than in trawls (V_43_ = 817, *P* < 0.001,) as well as for each estuary individually (Ashepoo; V_15_ = 103, *P* = 0.012; Dawho; V_13_ = 78, *P* = 0.021; Ashley; V_15_ = 106, *P* = 0.007, Figure 2). The mean ± SE difference between percent of males in pots and trawls was 20.1 ± 4.8 % overall, and 12.8 ± 6.3 %, 21.5 ± 10.5 %, and 26.7 ± 8.4 % for the Ashepoo, Dawho, and Ashley Rivers, respectively.

**Figure 2.**
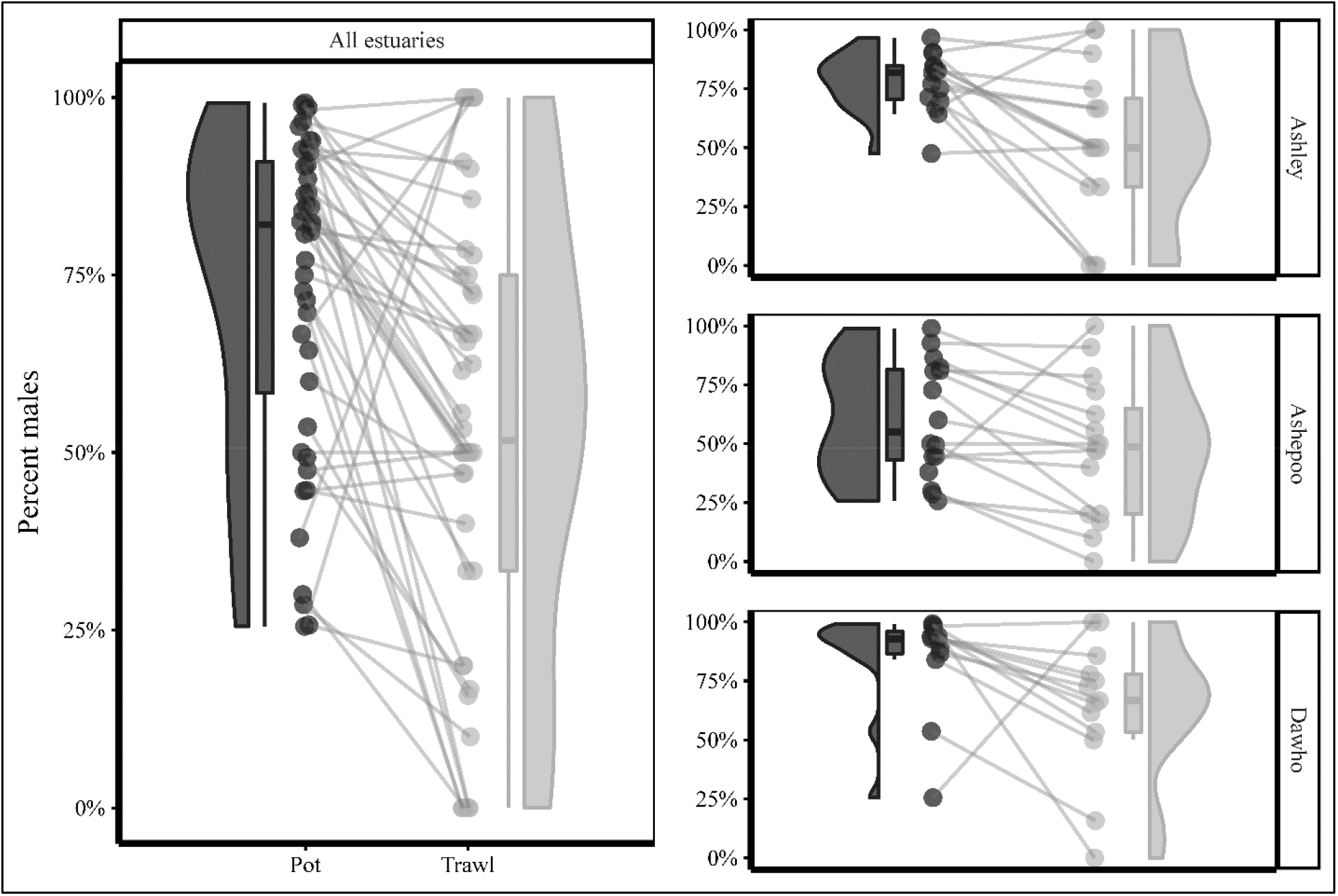
Density distribution plot, boxplot with median values and individual data points for percent male blue crabs collected in pot-based (dark grey) and trawl-based (light grey) fall sampling for all estuaries combined (left) and in three SC estuaries plotted separately (right). Points connected by lines indicate discrete sampling pairs. Significant (*P* <0.05) male bias was found for all estuaries combined and for each of the estuaries separately (see Results for details).

### [B] Within estuary year-round comparisons

For the year-round sampling in the Ashley River, overall, the percent of male crabs in pots was significantly higher than in trawls (V_59_ = 1509, *P* < 0.001). The overall mean ± SE % males was 66.8 ± 2.5 % and 44.1 ± 3.7 % for pots and trawls, respectively. Sex ratio bias towards males in pot sampling, however, varied by month. The mean percent males in pots was significantly greater than in trawls in June (V_17_ = 150, *P* < 0.001) and December (V_9_ = 41.5, *P* = 0.023), and marginally significant in April (V_9_ = 39, *P* = 0.055) and October (V_8_ = 32, *P* = 0.055). No significant differences were found for February (V_4_ = 1, *P* = 0.25), August (V_7_ = 14, *P* = 1.0) or November (V_5_ = 14, *P* = 0.125) (Figure 3). The mean percent of males ranged from 47.4 % to 79.7 % in pots and 27.6 % to 78.7 % in trawls.

**Figure 3.**
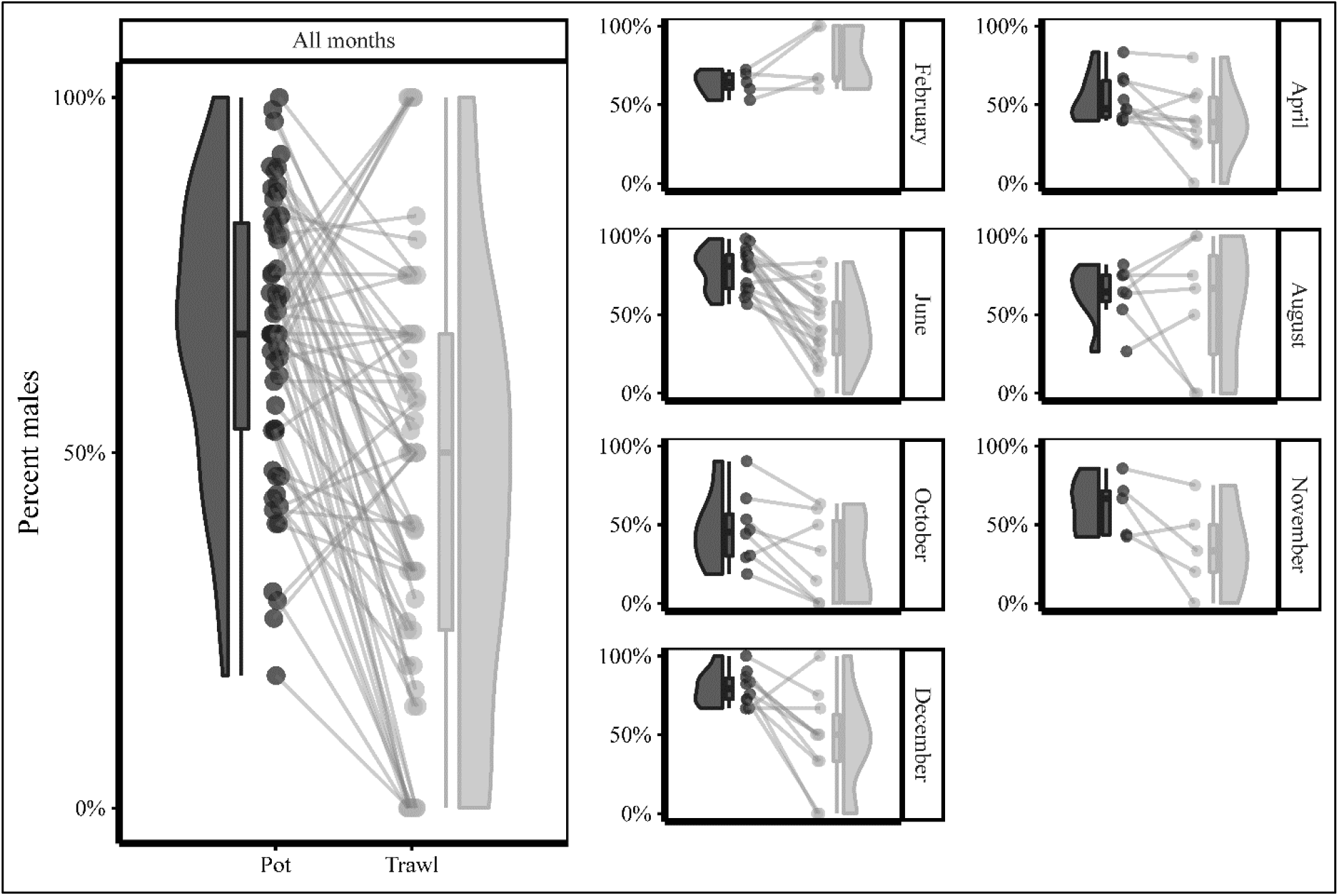
Density distribution plot, boxplot with median values and individual data points for percent male blue crabs collected in pot-based (dark grey) and trawl-based (light grey) year-round sampling in the Ashley River, SC, overall (left) and by month sampled (right). Points connected by lines indicate discrete sampling pairs. Significant (*P* <0.05) male bias was found for all months combined and for sampling conducted in June and December, while marginally significant (*P* = 0.055) male bias was found in April and October (see Results for details).

## [A] Discussion

The current study demonstrated that sampling of Atlantic Blue Crab in South Carolina using pot-based sampling approaches, while effective at collecting Atlantic Blue Crabs in substantial numbers, yielded sex ratios of catches that were significantly skewed towards male crabs relative to trawl-based sampling. Apparent disparities in sex selectivity of gear types could be due to a number of factors, including differences in the timing of sampling, Atlantic Blue Crab microhabitat distribution and behavior within those habitats, interactions with conspecifics and gear, and differential retention between gear types.

Segregation of Atlantic Blue Crab by sex is documented to occur at a large scale in the fall along the Atlantic coast, in part due to the migration of female crabs towards higher salinity waters to spawn (Hines et al. 1987). Ogburn and Habegger (2015) concluded that mature female crabs use estuaries as “corridors” during their seaward migration prior to spawning, which occurs largely on the continental shelf in the Georgia Bight. This behavior likely explains observed decreases in the proportion of female crabs in estuarine samples in the fall in both South Carolina (Archambault et al. 1990) and North Carolina (Ramach et al. 2009). Differences in the sex ratio in pot-based sampling relative to trawl-based sampling in the current study could have been due to such seasonal migratory patterns, as population demographics within the sampling space may have changed in the 14-day period between sampling events. Migration of female crabs out of the sampling area, however, seems an unlikely explanation for the male biased sex ratio in pots relative to trawls in the current study. If female crabs migrated out of the sampling areas within the 14-day window between sampling events, one might expect the sex ratio in December, when trawl sampling occurred, to be skewed towards male crabs. The finding of a statistically significant bias towards male crabs in pots relative to trawls in our analyses of sampling in the fall, irrespective of the number of days between sampling events, provides evidence that migration patterns do not fully explain the observed sex bias in pot sampling.

Segregation by sex has also been shown to occur at a smaller scale. Hines et al. (1987) reported segregation by sex in Atlantic Blue Crab for a sub-estuary of the Chesapeake Bay, where males are typically found in lower salinity creek heads and females tend to occur in higher salinity habitats. These authors also reported a slightly increased percent of male crabs in the composition of trawl samples in the late fall, with no mature females collected after October. An affinity of male crabs for lower salinity waters, or the fall migration of females seaward, may explain differences in sex ratios between gear types for the Dawho River site, given that sampling areas between the surveys did not overlap (∼1.2 km between survey areas) and that mean salinity in the fall was higher for the trawl survey (27.4 psu) than the potting survey (25.9 psu). The finding by Ramach et al. (2009) that segregation of sexes occurs in a small embayment, where salinity is not stratified suggests, however, that other environmental or habitat factors may also be at play. These other factors (*e*.*g*., vegetation and bathymetry, Cheng et al. 2022) may provide a mechanism for segregation to occur at the “microhabitat” scale and explain differences in sex ratios between gear types in some cases. Microhabitat scale differences could explain the sex bias towards male crabs in pots in the current study. Although the sampling areas of the two surveys in the Ashley and Ashepoo Rivers overlap within the watercourse, some differences in the exact location of sampling did occur. While pot placement for sampling typically focuses on locations closer to the shore, trawl sampling is generally conducted in the deeper channels. Male crabs may be selecting shallower water (Schweitzer and Withers 2009) and therefore be more susceptible to being captured in pots.

Although male and female Flower Crabs have been shown to be equally likely to enter pots (Smith and Sumpton 1989), we could find no empirical studies documenting the relative likelihood of male and female Atlantic Blue Crabs entering commercial crab pots. Female crabs may preferentially enter pots during certain times, however, particularly during periods of active mating when immature females nearing their pubertal (terminal) molt are attracted to mature male crabs for the purpose of copulation. This forms the basis of an important “peeler” crab fishery throughout the native geographic range of the Atlantic Blue Crab (Bishop et al. 1983; Sturdivant and Clark 2011), in which “peeler” pots are baited with large mature male crabs to support the supply of peeler crab stock for the soft-shell crab industry. The high abundance of sublegal-sized female crabs in pots in February and August relative to their low abundance in trawls in our sampling supports the premise that immature female crabs are likely to enter pots during the periods of highest mating activity in the Charleston area during the spring and fall (Archambault et al. 1990). This may be particularly important early in those seasons when larger mature males are less abundant (SCDNR, unpubl. data) and may explain why a male bias in pots was not observed in February and August in the current study.

In addition to the behavior of female Atlantic Blue Crabs potentially impacting the overall composition of pot-based samples, inefficiencies in the retention of females that enter the commercial-style pots used in our survey may have influenced the sex ratios. Guillory (1998) found that all male crabs in the 122-126 mm CW size range were retained in pots (38 mm mesh) without cull rings, whereas only 17% of females in this size range were retained. The author attributes this to the smaller carapace length, relative to carapace width, of females compared to males. Sturdivant and Clark (2011) found that commercial pots were inefficient at retaining legal male crabs, which were shown to enter and exit the lower compartment (kitchen) of pots with relative ease, suggesting that differential retention may influence sex ratios within pot samples in some cases. Once in the upper compartment (parlor) of pots, however, almost all male crabs were retained. These authors suggested that only crabs collected from the parlor sections of pots should be used for assessments of population dynamics. Because the location of crabs retained within pots (kitchen or parlor) could not be determined in the current study, comparisons of sex ratios within the parlor were not possible. More research is needed in order to understand sex-specific crab retention rates in commercial pots.

Management approaches for Atlantic Blue Crabs frequently rely on an understanding of sex-specific abundances within populations, often leading to restrictions on female harvest (Eggleston et al. 2009; Bunnell et al. 2010; Perry et al. 2022). Maintaining balanced sex-ratios within populations is important for their sustainability and to avoid potential impacts such as those related to male-focused harvest, specifically when the commercial fishery is pot-based (Carver et al. 2004; Ogburn et al. 2019). Accurate determination of sex ratios is also important for conducting rigorous Atlantic Blue Crab stock assessments that use sex-specific methods for management purposes (Miller et al. 2011; NCDMF 2018). Such management approaches should be based on assessments from sampling that accurately characterizes the population. The current study suggests that sampling of Atlantic Blue Crabs using trawl-based and pot-based sampling approaches may lead to inconsistent inferences about the demographics of the surveyed populations. Assessments could be further confounded by inter-seasonal variations in the direction and magnitude of sex biases due to changes in Atlantic Blue Crab behavior. Gear-specific differences in Atlantic Blue Crab sex ratios should be considered when evaluating sampling approaches for use in blue crab population assessments involving sex specific models.

## [A] Acknowledgements

Support for this project was generously provided from both the South Carolina Saltwater Recreational Fishing License Fund, Morgan Island Programmatic Income Fund, and state-appropriated funds managed by South Carolina Department of Natural Resources’ (SCDNR) Marine Resources Division based in Charleston, South Carolina, USA. The data used in this study would not be available without the hard work over multiple decades of staff working within both SCDNR’s Marine Resources Research Institute (MRRI) and the Office of Fisheries Management, and most recently in particular the efforts of past and current members of the MRRI’s Crustacean Research and Monitoring Section. There is no conflict of interest declared in this article.

## Notes

### Competing Interest Statement

The authors have declared no competing interest.

