## Supplementary Tables for "Evidence for a Male Bias in Atlantic Blue Crab Pot-Based Sampling"

**Supplementary Table 1. Sex specific catches (CPUE) of Atlantic Blue Crabs and the number of days between sampling events for paired trawl and pot sampling at three estuaries in fall sampling.**

| Year | Estuary | Sampling interval (d) | Female Trawl CPUE | Male Trawl CPUE | Total Trawl CPUE | Female Pot CPUE | Male Pot CPUE | Total Pot CPUE | Percent Males in Trawl | Percent Males in Pots |
|---|---|---|---|---|---|---|---|---|---|---|
| 1990 | Ashepoo | 6 | 3 | 5 | 8 | 3 | 14 | 17 | 62.5 | 82.4 |
| 1991 | Ashepoo | 7 | 6 | 22 | 28 | 18 | 77 | 95 | 78.6 | 81.1 |
| 1993 | Ashepoo | 6 | 1 | 10 | 11 | 9 | 114 | 123 | 90.9 | 92.7 |
| 1994 | Ashepoo | 5 | 5 | 13 | 18 | 1 | 95 | 96 | 72.2 | 99 |
| 1995 | Ashepoo | 14 | 9 | 8 | 17 | 46 | 37 | 83 | 47.1 | 44.6 |
| 1996 | Ashepoo | -9 | 8 | 2 | 10 | 6 | 16 | 22 | 20 | 72.7 |
| 1997 | Ashepoo | 10 | 3 | 3 | 6 | 5 | 21 | 26 | 50 | 80.8 |
| 1998 | Ashepoo | 8 | 2 | 0 | 2 | 14 | 6 | 20 | 0 | 30 |
| 1999 | Ashepoo | -9 | 4 | 5 | 9 | 6 | 38 | 44 | 55.6 | 86.4 |
| 2000 | Ashepoo | 4 | 9 | 8 | 17 | 16 | 24 | 40 | 47.1 | 60 |
| 2007 | Ashepoo | 11 | 5 | 1 | 6 | 36 | 35 | 71 | 16.7 | 49.3 |
| 2009 | Ashepoo | 12 | 18 | 2 | 20 | 30 | 12 | 42 | 10 | 28.6 |
| 2012 | Ashepoo | 6 | 4 | 1 | 5 | 26 | 9 | 35 | 20 | 25.7 |
| 2016 | Ashepoo | 8 | 0 | 1 | 1 | 62 | 38 | 100 | 100 | 38 |
| 2019 | Ashepoo | 10 | 6 | 4 | 10 | 26 | 21 | 47 | 40 | 44.7 |
| 2021 | Ashepoo | 11 | 3 | 3 | 6 | 1 | 1 | 2 | 50 | 50 |
| 1997 | Ashley | -8 | 3 | 0 | 3 | 10 | 23 | 33 | 0 | 69.7 |
| 1998 | Ashley | 3 | 1 | 2 | 3 | 8 | 24 | 32 | 66.7 | 75 |
| 1999 | Ashley | -8 | 1 | 9 | 10 | 3 | 86 | 89 | 90 | 96.6 |
| 2001 | Ashley | -1 | 2 | 2 | 4 | 21 | 19 | 40 | 50 | 47.5 |
| 2004 | Ashley | 10 | 0 | 2 | 2 | 4 | 38 | 42 | 100 | 90.5 |
| 2005 | Ashley | 9 | 4 | 4 | 8 | 12 | 54 | 66 | 50 | 81.8 |
| 2006 | Ashley | 8 | 3 | 3 | 6 | 8 | 44 | 52 | 50 | 84.6 |
| 2007 | Ashley | 14 | 1 | 2 | 3 | 11 | 37 | 48 | 66.7 | 77.1 |
| 2008 | Ashley | -13 | 1 | 1 | 2 | 5 | 28 | 33 | 50 | 84.8 |
| 2009 | Ashley | 7 | 1 | 0 | 1 | 26 | 47 | 73 | 0 | 64.4 |
| 2014 | Ashley | -1 | 1 | 0 | 1 | 3 | 14 | 17 | 0 | 82.4 |
| 2015 | Ashley | -8 | 2 | 1 | 3 | 6 | 56 | 62 | 33.3 | 90.3 |
| 2016 | Ashley | -9 | 1 | 3 | 4 | 7 | 33 | 40 | 75 | 82.5 |
| 2018 | Ashley | -9 | 2 | 1 | 3 | 12 | 30 | 42 | 33.3 | 71.4 |
| 2021 | Ashley | 9 | 0 | 1 | 1 | 4 | 8 | 12 | 100 | 66.7 |
| 1994 | Dawho | 1 | 3 | 8 | 11 | 22 | 142 | 164 | 72.7 | 86.6 |
| 1995 | Dawho | -6 | 21 | 40 | 61 | 11 | 85 | 96 | 65.6 | 88.5 |
| 1997 | Dawho | 12 | 4 | 14 | 18 | 3 | 46 | 49 | 77.8 | 93.9 |
| 1998 | Dawho | 14 | 16 | 3 | 19 | 45 | 52 | 97 | 15.8 | 53.6 |
| 2001 | Dawho | 9 | 0 | 1 | 1 | 35 | 12 | 47 | 100 | 25.5 |
| 2002 | Dawho | 14 | 5 | 8 | 13 | 1 | 69 | 70 | 61.5 | 98.6 |
| 2007 | Dawho | -9 | 7 | 8 | 15 | 3 | 70 | 73 | 53.3 | 95.9 |
| 2008 | Dawho | 4 | 1 | 3 | 4 | 4 | 52 | 56 | 75 | 92.9 |
| 2011 | Dawho | 13 | 0 | 1 | 1 | 3 | 165 | 168 | 100 | 98.2 |
| 2013 | Dawho | 11 | 2 | 4 | 6 | 5 | 78 | 83 | 66.7 | 94 |
| 2014 | Dawho | -9 | 1 | 1 | 2 | 8 | 42 | 50 | 50 | 84 |
| 2016 | Dawho | 9 | 1 | 0 | 1 | 1 | 133 | 134 | 0 | 99.3 |
| 2019 | Dawho | 11 | 1 | 6 | 7 | 3 | 36 | 39 | 85.7 | 92.3 |

**Supplementary Table 2. Sex specific catches (CPUE) of Atlantic Blue Crabs and the number of days between sampling events for paired trawl and pot sampling in year-round sampling in the Ashley River.**

| Year | Month | Sampling interval (d) | Female Trawl CPUE | Male Trawl CPUE | Total Trawl CPUE | Female Pot CPUE | Male Pot CPUE | Total Pot CPUE | Percent Males in Trawl | Percent Males in Pots |
|---|---|---|---|---|---|---|---|---|---|---|
| 2003 | April | -14 | 31 | 11 | 42 | 8 | 7 | 15 | 26.2 | 46.7 |
| 2003 | June | 14 | 6 | 1 | 7 | 6 | 25 | 31 | 14.3 | 80.6 |
| 2003 | October | 1 | 1 | 0 | 1 | 16 | 7 | 23 | 0 | 30.4 |
| 2004 | April | 2 | 25 | 16 | 41 | 21 | 19 | 40 | 39 | 47.5 |
| 2004 | June | -1 | 5 | 7 | 12 | 1 | 29 | 30 | 58.3 | 96.7 |
| 2004 | August | 3 | 5 | 5 | 10 | 22 | 8 | 30 | 50 | 26.7 |
| 2005 | February | 2 | 2 | 4 | 6 | 8 | 9 | 17 | 66.7 | 52.9 |
| 2005 | April | 2 | 3 | 2 | 5 | 18 | 13 | 31 | 40 | 41.9 |
| 2005 | June | 6 | 1 | 2 | 3 | 3 | 26 | 29 | 66.7 | 89.7 |
| 2005 | August | 14 | 1 | 2 | 3 | 11 | 19 | 30 | 66.7 | 63.3 |
| 2005 | December | -9 | 4 | 4 | 8 | 7 | 22 | 29 | 50 | 75.9 |
| 2006 | February | 4 | 2 | 3 | 5 | 12 | 18 | 30 | 60 | 60 |
| 2006 | April | -14 | 3 | 4 | 7 | 12 | 8 | 20 | 57.1 | 40 |
| 2006 | June | 7 | 1 | 1 | 2 | 10 | 20 | 30 | 50 | 66.7 |
| 2006 | October | 5 | 2 | 1 | 3 | 16 | 14 | 30 | 33.3 | 46.7 |
| 2006 | December | -8 | 3 | 3 | 6 | 5 | 25 | 30 | 50 | 83.3 |
| 2007 | June | -5 | 1 | 5 | 6 | 6 | 25 | 31 | 83.3 | 80.6 |
| 2007 | August | -12 | 1 | 0 | 1 | 14 | 16 | 30 | 0 | 53.3 |
| 2007 | December | -14 | 1 | 2 | 3 | 10 | 20 | 30 | 66.7 | 66.7 |
| 2008 | June | -6 | 4 | 1 | 5 | 4 | 26 | 30 | 20 | 86.7 |
| 2008 | December | 13 | 1 | 1 | 2 | 4 | 26 | 30 | 50 | 86.7 |
| 2009 | February | 13 | 0 | 1 | 1 | 5 | 13 | 18 | 100 | 72.2 |
| 2009 | April | 1 | 1 | 4 | 5 | 5 | 25 | 30 | 80 | 83.3 |
| 2009 | June | 7 | 2 | 1 | 3 | 13 | 17 | 30 | 33.3 | 56.7 |
| 2009 | December | -7 | 1 | 0 | 1 | 8 | 21 | 29 | 0 | 72.4 |
| 2010 | April | -1 | 3 | 1 | 4 | 5 | 10 | 15 | 25 | 66.7 |
| 2010 | June | -8 | 12 | 5 | 17 | 2 | 23 | 25 | 29.4 | 92 |
| 2011 | June | -6 | 17 | 11 | 28 | 13 | 22 | 35 | 39.3 | 62.9 |
| 2011 | August | 6 | 0 | 1 | 1 | 6 | 18 | 24 | 100 | 75 |
| 2011 | October | -1 | 1 | 0 | 1 | 19 | 15 | 34 | 0 | 44.1 |
| 2012 | April | 7 | 3 | 0 | 3 | 15 | 17 | 32 | 0 | 53.1 |
| 2012 | June | 14 | 3 | 2 | 5 | 6 | 24 | 30 | 40 | 80 |
| 2012 | October | -2 | 1 | 1 | 2 | 17 | 7 | 24 | 50 | 29.2 |
| 2013 | June | 5 | 5 | 1 | 6 | 5 | 25 | 30 | 16.7 | 83.3 |
| 2014 | June | -3 | 1 | 3 | 4 | 9 | 21 | 30 | 75 | 70 |
| 2014 | December | 1 | 1 | 0 | 1 | 3 | 14 | 17 | 0 | 82.4 |
| 2015 | June | 13 | 1 | 0 | 1 | 7 | 14 | 21 | 0 | 66.7 |
| 2015 | October | -3 | 7 | 12 | 19 | 9 | 84 | 93 | 63.2 | 90.3 |
| 2015 | December | 8 | 2 | 1 | 3 | 6 | 55 | 61 | 33.3 | 90.2 |
| 2016 | April | -2 | 2 | 1 | 3 | 21 | 14 | 35 | 33.3 | 40 |
| 2016 | June | 2 | 6 | 2 | 8 | 18 | 28 | 46 | 25 | 60.9 |
| 2016 | August | 2 | 1 | 0 | 1 | 4 | 18 | 22 | 0 | 81.8 |
| 2016 | October | 1 | 6 | 1 | 7 | 7 | 8 | 15 | 14.3 | 53.3 |
| 2016 | November | -5 | 1 | 1 | 2 | 19 | 14 | 33 | 50 | 42.4 |
| 2016 | December | 9 | 1 | 3 | 4 | 0 | 10 | 10 | 75 | 100 |
| 2017 | February | -14 | 1 | 2 | 3 | 7 | 16 | 23 | 66.7 | 69.6 |
| 2017 | June | 8 | 2 | 1 | 3 | 6 | 41 | 47 | 33.3 | 87.2 |
| 2017 | August | 12 | 0 | 1 | 1 | 16 | 29 | 45 | 100 | 64.4 |
| 2018 | June | 3 | 15 | 17 | 32 | 1 | 57 | 58 | 53.1 | 98.3 |
| 2018 | August | 11 | 1 | 3 | 4 | 21 | 64 | 85 | 75 | 75.3 |
| 2018 | October | -4 | 1 | 0 | 1 | 35 | 8 | 43 | 0 | 18.6 |
| 2018 | November | 1 | 4 | 1 | 5 | 35 | 27 | 62 | 20 | 43.5 |
| 2018 | December | 9 | 2 | 1 | 3 | 11 | 29 | 40 | 33.3 | 72.5 |
| 2019 | February | 7 | 0 | 4 | 4 | 10 | 18 | 28 | 100 | 64.3 |
| 2019 | April | 2 | 5 | 6 | 11 | 22 | 41 | 63 | 54.5 | 65.1 |
| 2019 | June | -7 | 11 | 15 | 26 | 5 | 36 | 41 | 57.7 | 87.8 |
| 2019 | November | 12 | 1 | 0 | 1 | 5 | 10 | 15 | 0 | 66.7 |
| 2020 | October | 7 | 2 | 3 | 5 | 8 | 16 | 24 | 60 | 66.7 |
| 2020 | November | 2 | 1 | 3 | 4 | 4 | 24 | 28 | 75 | 85.7 |
| 2021 | November | 3 | 2 | 1 | 3 | 2 | 5 | 7 | 33.3 | 71.4 |
| 2021 | December | -9 | 0 | 1 | 1 | 4 | 8 | 12 | 100 | 66.7 |

**Supplementary Table 3. Summary data and results of statistical analysis for differences in percent male blue crabs collected in pot-based and trawl-based sampling for the maximum number of days between paired trawl and pot sampling events.**

| | | Percent male crabs in pot sampling | | | | Percent male crabs in trawl sampling | | | | % males in pots - % males in trawls | | | | Wilcoxon (two-tailed) test results | | | | | | |
|---|---|---|---|---|---|---|---|---|---|---|---|---|---|---|---|---|---|---|---|---|
| Maximum Sampling Interval (days) | Survey | Mean | sd | SE | Median | Mean | sd | SE | Median | Mean | sd | SE | Median | V | p-value | (pseudo) median | n | W | W- | W+ |
| 14 | Fall | 73.5 | 22.3 | 3.4 | 82.1 | 53.4 | 30.7 | 4.6 | 51.7 | 20.1 | 31.6 | 4.8 | 19.2 | 817 | ≤0.0001 | 20.6 | 43 | 946 | 129 | 817 |
| 10 | Fall | 74 | 21.6 | 3.8 | 82.1 | 53.2 | 32 | 5.7 | 51.7 | 20.9 | 36.1 | 6.4 | 21.4 | 439 | 0.0007 | 21.3 | 32 | 528 | 89 | 439 |
| 7 | Fall | 75.2 | 20.8 | 5.8 | 82.4 | 53.9 | 29.6 | 8.2 | 65.6 | 21.3 | 24.9 | 6.9 | 13.9 | 88 | 0.0012 | 15.4 | 13 | 91 | 3 | 88 |
| 5 | Fall | 77.6 | 18.3 | 6.9 | 82.4 | 54.8 | 26.6 | 10.1 | 66.7 | 22.8 | 27.7 | 10.5 | 13.9 | 27 | 0.0313 | 15.6 | 7 | 28 | 1 | 27 |
| 14 | Year-round | 66.8 | 19.1 | 2.5 | 66.7 | 44.1 | 29.4 | 3.8 | 50 | 22.7 | 30.7 | 3.9 | 23.5 | 1509 | ≤0.0001 | 23.5 | 59 | 1770 | 261 | 1509 |
| 10 | Year-round | 66.9 | 20.7 | 3 | 66.7 | 43.2 | 28 | 4.1 | 40 | 23.7 | 29.4 | 4.3 | 23.6 | 946 | ≤0.0001 | 24.3 | 46 | 1081 | 135 | 946 |
| 7 | Year-round | 63.5 | 20.7 | 3.3 | 65.1 | 41.7 | 28.6 | 4.6 | 40 | 21.8 | 29.2 | 4.7 | 23 | 638 | ≤0.0001 | 21.9 | 38 | 741 | 103 | 638 |
| 5 | Year-round | 60.5 | 22.6 | 4.3 | 60.4 | 39.3 | 26.4 | 5 | 39.5 | 21.2 | 28 | 5.3 | 16 | 328 | 0.0004 | 20.7 | 27 | 378 | 50 | 328 |
| 3 | Year-round | 61.9 | 22.4 | 4.8 | 63 | 38.9 | 26 | 5.5 | 39.5 | 23 | 28.5 | 6.1 | 25.4 | 223 | 0.0009 | 22.5 | 22 | 253 | 30 | 223 |
| 1 | Year-round | 62.6 | 23.4 | 8.3 | 60 | 24.7 | 29.6 | 10.5 | 17.1 | 37.9 | 22.3 | 7.9 | 38.7 | 36 | 0.0078 | 37.8 | 8 | 36 | 0 | 36 |

**Supplementary Table 4. Summary data and results of statistical analysis for differences in percent male blue crabs collected in pot-based and trawl-based sampling in the fall in three estuaries and year-round in a single estuary. The maximum interval between trawl and pot sampling was 14 days for the analysis.**

| | | Percent male crabs in pot sampling | | | | Percent male crabs in trawl sampling | | | | % males in pots - % males in trawls | | | | Wilcoxon (two-tailed) test results | | | | | | |
|---|---|---|---|---|---|---|---|---|---|---|---|---|---|---|---|---|---|---|---|---|
| | Survey | Mean | sd | SE | Median | Mean | sd | SE | Median | Mean | sd | SE | Median | V | p-value | (pseudo) median | n | W | W- | W+ |
| Overall | Fall | 73.5 | 22.3 | 3.4 | 82.1 | 53.4 | 30.7 | 4.6 | 51.7 | 20.1 | 31.6 | 4.8 | 19.2 | 817 | ≤0.0001 | 20.6 | 43 | 946 | 129 | 817 |
| Ashepoo | Fall | 60.4 | 24.6 | 6.1 | 55 | 47.5 | 29.1 | 7.3 | 48.5 | 12.8 | 25.2 | 6.3 | 15.8 | 103 | 0.0125 | 16.3 | 15 | 120 | 17 | 103 |
| Dawho | Fall | 84.9 | 21.4 | 5.9 | 92.9 | 63.4 | 29.1 | 8.1 | 66.7 | 21.5 | 37.9 | 10.5 | 23 | 78 | 0.0215 | 23 | 13 | 91 | 13 | 78 |
| Ashley | Fall | 77.7 | 12.4 | 3.2 | 81.8 | 51 | 33.6 | 8.7 | 50 | 26.7 | 32.4 | 8.4 | 31.8 | 106 | 0.0067 | 24.4 | 15 | 120 | 14 | 106 |
| Overall | Year-round | 66.8 | 19.1 | 2.5 | 66.7 | 44.1 | 29.4 | 3.7 | 50 | 22.7 | 30.7 | 3.9 | 23.5 | 1509 | ≤0.0001 | 23.5 | 59 | 1770 | 261 | 1509 |
| February | Year-round | 63.8 | 7.7 | 3.4 | 64.3 | 78.7 | 19.7 | 8.8 | 66.7 | -14.9 | 16.9 | 7.5 | -13.7 | 1 | 0.2500 | -18.6 | 4 | 10 | 9 | 1 |
| April | Year-round | 53.8 | 14.9 | 5 | 47.5 | 39.5 | 22.8 | 7.6 | 39 | 14.3 | 21.4 | 7.1 | 8.5 | 39 | 0.0547 | 11.2 | 9 | 45 | 6 | 39 |
| June | Year-round | 79.2 | 12.9 | 3.1 | 80.6 | 40.9 | 22.8 | 5.5 | 39.3 | 38.3 | 23.5 | 5.7 | 38.3 | 150 | ≤0.0001 | 39 | 17 | 153 | 3 | 150 |
| August | Year-round | 62.8 | 18.6 | 7 | 64.4 | 56 | 42.1 | 15.9 | 66.7 | 6.9 | 44.1 | 16.7 | -3.3 | 14 | 1.0000 | -0.6 | 7 | 28 | 14 | 14 |
| October | Year-round | 47.4 | 23 | 8.1 | 45.4 | 27.6 | 27.5 | 9.7 | 23.8 | 19.8 | 20.7 | 7.3 | 22.9 | 32 | 0.0547 | 22.4 | 8 | 36 | 4 | 32 |
| November | Year-round | 62 | 18.7 | 8.4 | 66.7 | 35.7 | 28.6 | 12.8 | 33.3 | 26.3 | 28.1 | 12.6 | 23.5 | 14 | 0.1250 | 24.4 | 5 | 15 | 1 | 14 |
| December | Year-round | 79.7 | 10.8 | 3.4 | 79.1 | 45.8 | 31.2 | 9.9 | 50 | 33.8 | 33.7 | 10.6 | 35 | 41.5 | 0.0234 | 37.3 | 9 | 45 | 3.5 | 41.5 |
